# A Proximity-Dependent Biosensor System for Visualizing Cell-Cell Interactions Induced by Therapeutic Antibodies

**DOI:** 10.1101/2022.05.04.490615

**Authors:** Yu Tang, Yanguang Cao

## Abstract

**Background and Purpose:** Despite the promising results of therapeutic antibodies in engaging the immune system to eliminate malignant cells, many aspects of the complex interplay between immune cells and cancer cells during antibody therapy remain incompletely understood. This study aimed to develop a biosensor system that can evaluate direct cell-cell contact and interactions between immune effector and target cells induced by therapeutic antibodies in physiologically relevant environments.

**Experimental Approach:** The system uses two structural complementary luciferase units (SmBit and LgBit) expressed on the respective membranes of immune effector and target cells. Upon cell-cell contact, the two subunits form active NanoLuc, generating a luminescent signal, allowing for real-time monitoring of cell-cell interactions and quantitatively assessing the pharmacological effects of therapeutic antibodies.

**Key Results:** We optimized the system to ensure selectivity by adjusting the spacer lengths between two luciferase units to minimize interference from nonspecific intercellular contact. The system was able to quantitatively and longitudinally monitor cell-cell interactions between NK and target cells induced by rituximab and between T and target cells induced by blinatumomab in a three-dimensional cell culture system. The observation that NK cells exhibited faster interactions with target cells compared to T cells is intriguing and suggests potential differences in the mechanisms or kinetics of cell-cell interactions between different types of effector cells and tumor cells.

**Conclusions and Implications:** The biosensor system has the potential for broad applications to optimize antibody pharmacology and efficacy in various therapeutic areas through a deeper understanding of antibody-mediated cell-cell interactions.

**What is already known:**

- Therapeutic antibodies can activate the immune system to elicit cytotoxicity through inducing cell-cell interactions between immune and tumor cells.
- Approaches for longitudinally evaluating cell-cell interaction induced by therapeutic antibodies remain limited.

**What does this study add:**

- This study develops a biosensor system for detecting cell-cell interactions induced by therapeutic antibodies in a longitudinal manner.
- The system is able to monitor cell-cell interactions between NK and target cells, as well as between T and target cells, in a three-dimensional cell culture system.

**What is the clinical significance:**

- The biosensor system has the potential for broad applications in the field of antibody pharmacology by providing a deeper understanding of antibody-mediated cell-cell interactions and their dynamics.

## Introduction

Therapeutic antibodies can engage innate and adaptive (T) immune cells to eliminate pathogenic or malignant cells (Redman, Hill, AlDeghaither & Weiner, 2015). Immune cell activation and intercellular interaction between immune effector and target cells are key to the pharmacological effects of many antibody-based therapeutic approaches, including but not limited to checkpoint blockades, circulating cytokine neutralizers, bispecific T cell engagers, and virus neutralizers (Tang & Cao, 2021; Waldman, Fritz & Lenardo, 2020). Characterizing antibody-induced cell-cell interactions between immune effector and target cells is therefore crucial to understanding antibody pharmacological effect and pharmacodynamics (PD), provide insights into the mechanisms of immune evasion and treatment resistance, and inform strategies for optimizing antibody therapeutics in various therapeutic areas (Bechtel, Reyes-Robles, Fadeyi & Oslund, 2021; Shelton, Nguyen, Barbie & Kamm, 2021).

Despite the fact that understanding the dynamics of cell-cell interactions and the formation of immunological synapse upon cell-cell interactions at the cell population level is critical for optimizing antibody-based therapeutics, to date, investigations in this area have focused primarily on molecular-scale interactions (Dustin, 2014a; Dustin, 2014b), leaving gaps in our understanding of the intercellular steps at the cell population level involved in effective immune cell activation and target cell killing. Cell population-level interactions are complex and involve dynamic and coordinated responses of multiple cells within a population (Armingol, Officer, Harismendy & Lewis, 2021). These interactions can be influenced by various factors (Nishida-Aoki & Gujral, 2019), such as cell density, spatial arrangement, and environmental cues, which can impact the overall effectiveness of immune cell activation and target cell killing mediated by therapeutic antibodies.

In pathological environments, such as tumor microenvironments, environmental factors can impair the functions of effector cells, including their ability to find and engage target cells (Baginska et al., 2013). These factors work together to create a niche that restricts effector cell infiltration, motility, adhesion, and effector functionality (Zhang et al., 2019). Thus, characterizing cell-cell interactions at the population level in physiologically relevant environments can provide valuable insights into the mechanisms of immune evasion and treatment resistance, and inform strategies for optimizing antibody-based therapeutics.

Here we constructed a proximity-based biosensor system to detect stable intercellular contact and interaction. The biosensor system described is designed to detect stable intercellular contact and interaction between immune effector and target cells using two structurally complementary luciferase subunits. Briefly, two structurally complementary luciferase subunits in the NanoBiT® system (Dixon et al., 2016), Large_BiT and Small_BiT, upon transfection, were respectively expressed on the surfaces of immune effector and tumor cells. Upon cell-cell contact and interaction, the proximity between tumor and immune cells allows the binding of Large_BiT and Small_BiT, forming active luciferases that emit strong luminescence upon substrate stimulus (Dixon et al., 2016). The system was further applied to detect NK-tumor cell interactions induced by an anti-CD20 antibody rituximab and T-tumor cell interactions induced by a bispecific T cell engaging antibody blinatumomab in a three-dimensional (3D) cell culture system. The biosensor system offers a promising approach to monitor and evaluate cell-cell interactions in relevant physiological conditions and to understand the factors that impede effective intercellular interactions induced by therapeutic antibodies.

## Materials and Methods

### Plasmid Construction

The pDisplay vector was used as a eukaryotic expression vector. It contains a murine Ig k chain leader sequence and a platelet-derived growth factor receptor transmembrane domain (PDGFR-TM) that are located at the N-terminus or C-terminus of the inserted gene. These elements help to direct and anchor the fusion protein to the cell membrane. Additionally, the vector contains HA and myc epitopes on both sides of the expressed recombinant proteins, which enable the detection of fusion peptides (Yang et al., 2007). Figure 1 depicts the schematic representation of the pDisplay vector and its key components.

**Figure 1.**
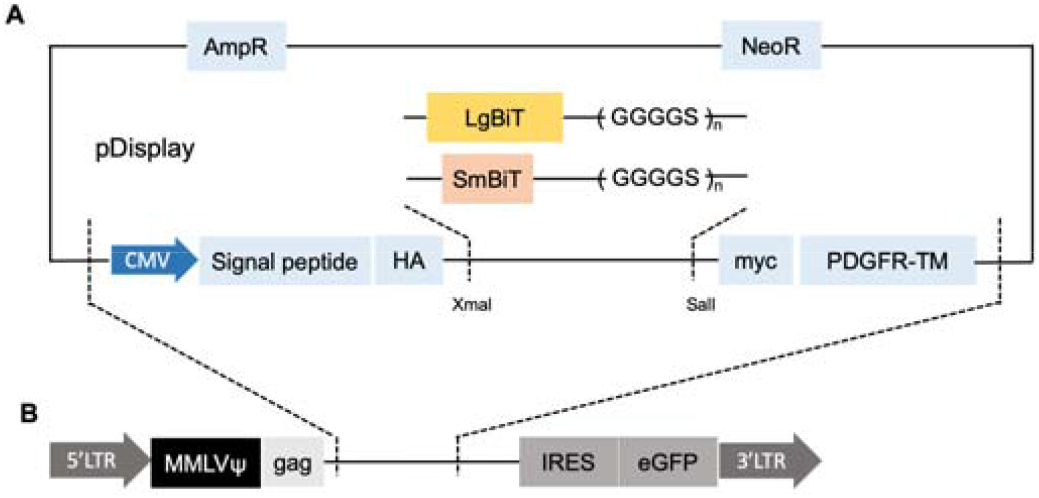
Construction of the biosensor system plasmid. (**A**) LgBiT or SmBiT constructed in pDisplay vector. (**B**) The CMV-LgBiT-(GGGGS)n-PDGFR-TM gene was cloned into PBMN-I-eGFP.

The sequences of SmBiT and LgBiT were kindly provided by Promega. Spacers were added to the C terminus of LgBiT or SmBiT to yield different distances between split luciferases and cell membranes. The LgBiT-(GGGGS)_n_ or SmBiT-(GGGGS)_n_ sequences were inserted into the pDisplay plasmid at XmaI and SaLI restriction sites. The CMV-LgBiT-(GGGGS)_n_-PDGFR-TM gene was cloned into PBMN-I-eGFP to produce recombinant retrovirus. PBMN-I-GFP was a gift from Garry Nolan (Addgene plasmid # 1736)

### Cell Culture and Transductions

HeLa cells were cultured in DMEM with 10% FBS (Invitrogen), 100 U/mL penicillin, and 100 mg/mL streptomycin (Gibco). Phoenix-AMPHO cells (ATCC^®^ CRL-3212™) were cultured in RPMI 1640 medium supplemented with 10% FBS (Invitrogen), 100 U/mL penicillin, and 100 mg/mL streptomycin (Gibco). Human Natural Killer cell line (NK-CD16; ATCC^®^ PTA-6967) was cultured in the Alpha-MEM medium. Cells were maintained in a humidified incubator at 37 °C.

SmBiT or LgBiT fusion genes expressed from pDisplay™ were introduced to HeLa cells via Lipofectamine 3000 (Invitrogen). NK-CD16 cells were transduced based on the protocol described before (Miah & Campbell, 2010). Briefly, the Phoenix-AMPHO cells were transfected with the engineered vector (PBMN-IRES-EGFP-SmBiT) via Lipofectamine 3000 to produce recombinant retrovirus. The retroviral supernatant was collected from transfected Phoenix-AMPHO cells for NK-CD16 transduction. NK-CD16 cells with positive SmBiT expression were selected via cell sorting by probing HA tag (anti-HA antibody [Invitrogen #26183, 1:500]; Anti-mouse secondary antibody Alexa Fluor 647 [Invitrogen #A-21235, 1:200]), probing CD16 (anti-CD16 antibody [Abcam #246222, 1:600]; Anti-rabbit secondary antibody Cy3 [Abcam #6939, 1:200]), and probing eGFP simultaneously.

### SmBiT/LgBiT Expressions on HeLa Cells

Transfected cells were harvested at 96 hour post-transfection, suspended in Opti-MEM, and transferred to a 96-well plate (1×10^5^ cells per well). HiBiT control protein, a 11 amino acid peptide with a higher affinity to LgBiT than SmBiT (0.7 nM vs. 190 μM), was used to detect LgBiT expression. NanoLuc substrate (furimazine) solution (1:50 diluted) containing 40 nM HiBiT control protein was added to the 96-well plate (20 μL per well). The luminescence was measured at 460 nm by Cytation 3 equipped with 460/40 nm bandpass filter.

The membranous and cytosolic LgBiT and SmBiT proteins were extracted from LgBiT-positive, SmBiT-positive, and HeLa WT cells using Mem-PER Plus membrane protein extraction kit. SmBiT expression was detected by LgBiT proteins. Membranous and cytosolic protein solutions were added to 96-well plates after adjusting the protein level. HiBiT-containing (40 nM) or LgBiT-containing (1:100 diluted) furimazine solution to detect LgBiT or SmBiT proteins.

To evaluate the binding function of the engineered SmBiT and LgBiT proteins at absence of cellular spatial limitations, HeLa cells were co-transfected or separately transfected by pDisplay-SmBiT and pDisplay-LgBiT genes via Lipofectamine 3000 (Invitrogen). Transfected cells were harvested at 96 hour post-transfection, suspended in Opti-MEM, and transferred to a 96-well plate (1×10^5^ cells per well). Furimazine solution was added to each well (1:50 diluted, 20 μL per well) 5-min before bioluminescence measurement with Cytation 3.

To test engineered SmBiT-LgBiT protein binding function with the cellular spatial limitations, HeLa cells separately expressed pDisplay-SmBiT or pDisplay-LgBiT 1:1 mixed, transferred to a 96-well plate, and briefly centrifuged at 200g before adding the furimazine solutions. The luminescence was measured at 460 nm by Cytation 3.

### SmBiT Expression on NK Cells and LgBiT expression on CD20^+^ HeLa Cells

SmBiT expression on NK-CD16-SmBiT cells was assessed in addition to HA^+^ sorting. NK-CD16-SmBiT cells were transferred to a 96-well plate (1×10^5^ cells per well). HeLa transfected with pDisplay-SmBiT genes (SmBiT-0L, 1L, 3L, and 12L) via Lipofectamine 3000 (Invitrogen) worked as positive controls. HeLa WT and NK-CD16 WT cells were used as negative controls. LgBiT-containing (1:100 diluted) furimazine solution was added to SmBiT^+^ cells to detect SmBiT expression levels. The S/N ratios were calculated as below:

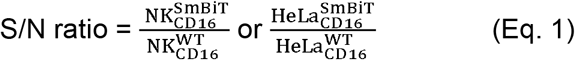

CD20 gene (Addgene plasmid # 1890) and pDisplay-LgBiT genes (LgBiT-0L, 1L, 3L, and 12L) were introduced to HeLa cells via Lipofectamine 3000 (Invitrogen). Transfected cells were harvested at 96 hour post-transfection. To determine CD20 expressions, transfected and WT Hela cells were washed with PBS, then incubated with FITC anti-CD20 antibody solution (BD B556632, 1:300 dilution) for 30 min. Each replicate included 1×10^5^ cells. After washing with PBS, FITC signal was quantified using Cytation 3 fluorescence monochromator at λex/em = 485/528 nm with a gain of 100.

### Antibody-induced effector and target cell-cell interactions in suspension

The ability of the biosensor system to detect antibody-induced cell clustering was validated in HeLa-EGFR:cetuximab:NK system. HeLa cells were transfected by pDisplay-LgBiT-12L cells via Lipofectamine 3000 (Invitrogen). NK-CD16-SmBiT-12L cells were mixed with pDisplay-LgBiT-12L-HeLa cells at E:T ratios of 10 or 5 (1×10^5^: 1×10^4^ or 5×10^4^: 1×10^4^ cells in 100 μL medium per well), with or without cetuximab (MedChemExpress) at concentration of 10 nM. NK-CD16 WT cells were mixed with pDisplay-LgBiT-12L-HeLa cells at the same conditions. After 30-min incubation, the luminescence was measured at 460 nm by Cytation 3.

### Imaging cell-cell interaction in a 3D cell culture system

At 96 hour post-transfection, CD20^+^LgBiT^+^ HeLa cells were harvested suspended in a 1:1 mixture of Matrigel (BD Bioscience). Matrigel-encapsulated cells were seeded in 24-well plates (20 μL in each well) and solidified at 37 °C for 30 min. In the 3D cell culture imaging studies, NK-CD16 cells with or without SmBiT expression were added to solidify 3D cell cultures with or without rituximab. The images were acquired at 30-min after substrate supplement (furimazine, 1:100 dilution) using an IVIS optical imaging system (Caliper Life Sciences) with an electron multiplying charge-coupled device camera. Acquired images were processed and quantified using Living image 4.5.2 (Caliper Life Sciences).

To compare the sensitivities in detecting rituximab-induced cell clustering between LgBiT-3L+SmBiT-12L and LgBiT-12L+SmBiT-12L, corrected bioluminescent signals were calculated as below:

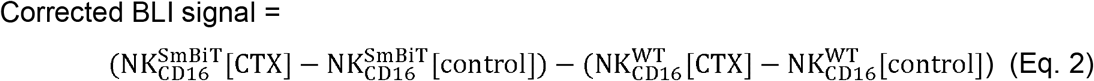

The peak corrected signals of each spacer combinations (LgBiT-0L+SmBiT-0L, LgBiT-1L+SmBiT-1L, LgBiT-3L+SmBiT-3L, LgBiT-0L+SmBiT-12L, LgBiT-1L+SmBiT-12L, LgBiT-3L+SmBiT-12L, and LgBiT-12L+SmBiT-12L) were calculated and normalized by control group signals:

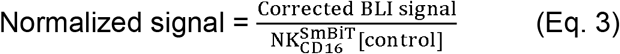

## Results

### The proximity-based biosensor system

The proximity-based biosensor system was engineered from NanoBiT, a complementary structural system composed of a Large BiT (LgBiT, 18 kDa) and a Small BiT (SmBiT, 11 amino acid peptide), which come together to form an active NanoLuc and generate bright luminescent signals. LgBiT and SmBiT were respectively expressed on target and effector cell membranes. Upon the LgBiT+ and SmBiT+ cell-cell contact, the two complementary parts on opposing cell surfaces come together and emit bright luminescent signals upon the addition of NanoLuc substrate.

The molecular bonds between LgBiT and SmBiT did not noticeably interfere with the natural intercellular interaction between effector and target cells, due to their low binding affinity (190 μM). To optimize the biosensor system, the spacer length between LgBiT and SmBiT was adjusted to ensure high specificity and sensitivity in detecting stable intercellular contact and immunological synapse formation (Figure 2A). Spacer-fused SmBiT biosensor components were constructed in plasmid DNA and viral vector forms. We tested the biosensor system with a wide range of combined spacer lengths from 0 – 48 nm (Table 1), and the optimized spacer lengths were selected for given pair of effector and target cells with the highest S/N ratios (Figure 2B, upper panel). When combined spacer lengths were too short, the sensitivity of detection by the biosensor system decreased (Figure 2B, middle panel). When combined spacer lengths were too long, the biosensor system would show high noise (i.e., background signal) with reduced selectivity for detecting stable cell-cell interaction (Figure 2B, lower panel).

**Figure 2.**
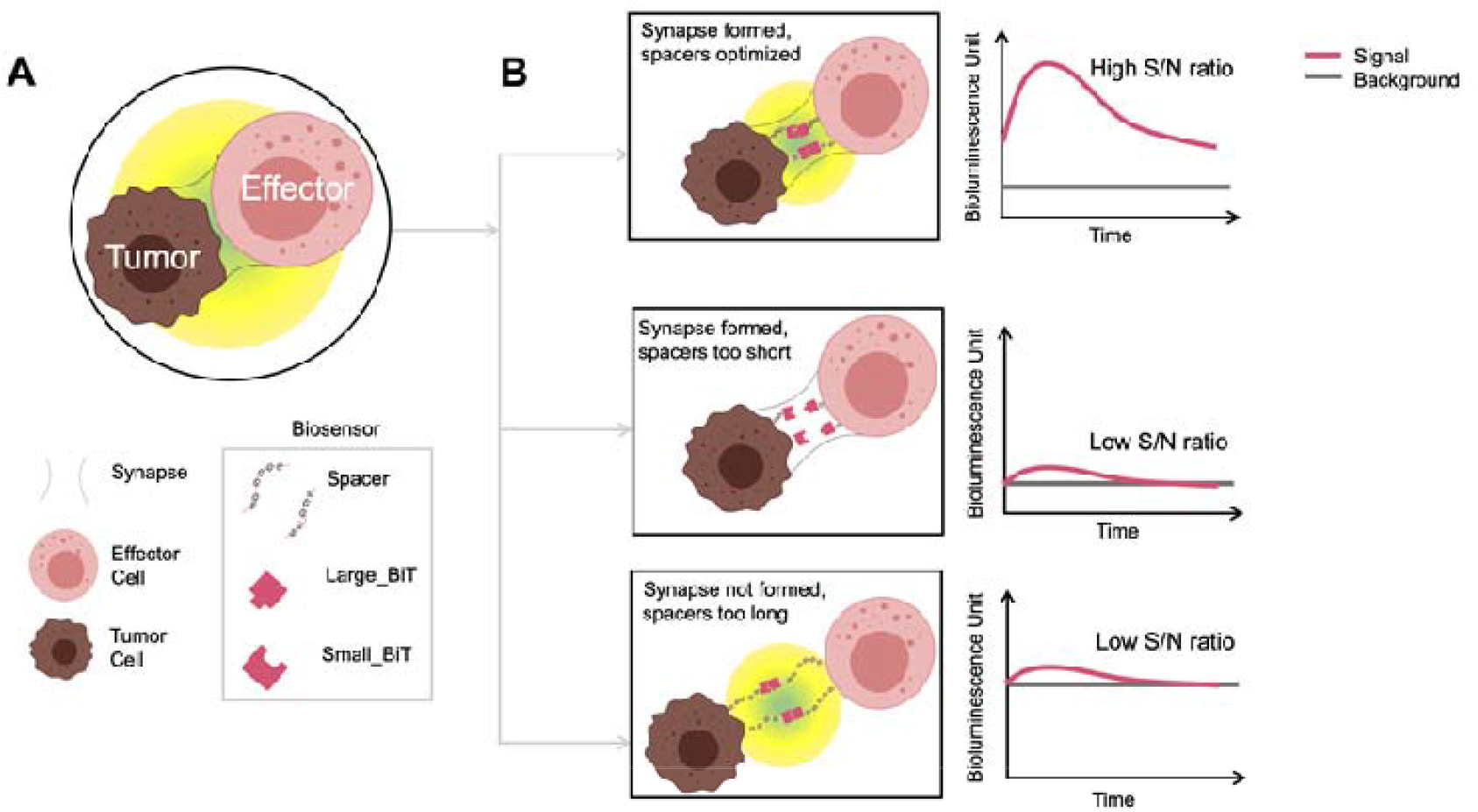
The design of the proximity-based luminescent biosensor system for detecting interactions between immune effector and target cells with optimized selectivity and sensitivity. (**A**) The schematic of the biosensor system. Briefly, two structurally complementary luciferase subunits in the NanoBiT® system, Large_BiT and Small_BiT, are separately anchored to the surfaces of immune and tumor cells by the spacers with various lengths. Once immunological synapses are formed, the proximity between tumor and immune cells allows integration of Large_BiT and Small_BiT, forming active luciferases that emit strong luminescence upon stimulus. The spacer used in constructing the biosensor system is (GGGGS)n, which was selected for its flexibility and linearity. The length of the GGGGS spacer is 1.9 nm per unit. (**B**) Optimizing the sensitivity and selectivity of the biosensor system by adjusting the combined spacer lengths. The biosensor system subunits with different spacer lengths facilitate the optimization of the signal-to-noise (S/N) ratios. When combined spacer lengths were too short, the sensitivity of detection by the biosensor system will decrease (middle panel). When combined spacer lengths were too long, the biosensor system will create high background signal and reduce the selectivity in detecting the formation of immunological synapses (lower panel).

**Table 1.** Construction of the biosensor system with different lengths of spacers.

| **LgBiT** |  | **SmBiT** |  |
| --- | --- | --- | --- |
| Biosensor subunit structure | Length (nm) | Biosensor subunit structure | Length (nm) |
| N'– LgBiT – C' | 0 | N'– SmBiT – C' | 0 |
| N'–LgBiT –(GGGGS)<sub>1</sub> –C' | 2 | N'–SmBiT –(GGGGS)<sub>1</sub> –C' | 2 |
| N'–LgBiT –(GGGGS)<sub>3</sub> –C' | 6 | N'–SmBiT –(GGGGS)<sub>3</sub> –C' | 6 |
| N'–LgBiT –(GGGGS)<sub>12</sub> –C' | 24 | N'–SmBiT –(GGGGS)<sub>12</sub> –C' | 24 |

### Engineered the biosensor system with high sensitivity

We characterized the expressions and functions of the biosensor system in HeLa cells. As Figure 3A shows, the LgBiT and SmBiT expressions in HeLa cells were significant compared to background (p < 0.0001). Both LgBiT and SmBiT were detectable in cytosols (Figure 3B), but their expressions on the cell membrane were much higher than in cytosols (p = 0.0006). The high membranous expression of the biosensor system suggests its potential for use in detecting intercellular interactions (Figure 3B).

**Figure 3.**
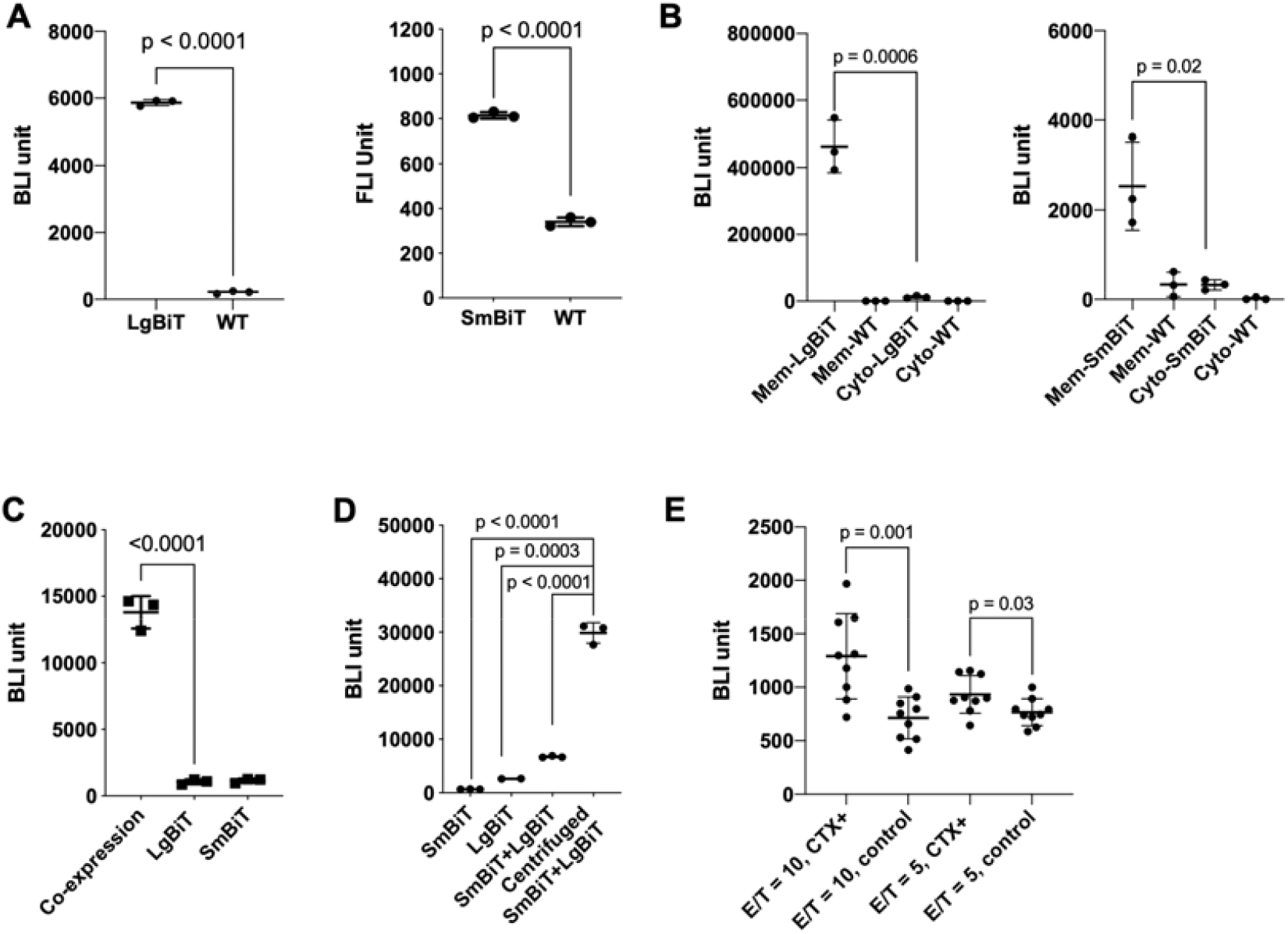
The biosensor system detected cell-cell interactions in a proximity-dependent manner. (**A**) LgBiT and SmBiT were successfully expressed in HeLa cells as indicated by HiBiT probing (Left panel) or HA tag probing (Right panel). (**B**) LgBiT and SmBiT were majorly expressed on cell membranes. Left panel: Compared to HeLa WT cells, LgBiT expression on membrane and in cytosol were significantly elevated. Membranous LgBiT expression was substantially higher than cytosolic expression. Right panel: Compared to HeLa WT cells, SmBiT expression on membrane and in cytosol were significantly elevated. Membranous SmBiT expression was substantially higher than cytosolic expression. (**C**) Co-expressed LgBiT and SmBiT emitted strong luminescent signals compared to LgBiT-or SmBiT-positive cells alone. (**D**) Biosensor system detected cell contacts that elevated by centrifugation. Compared to centrifuged LgBiT-or SmBiT-positive cells alone and uncentrifuged LgBiT:SmBiT cell mixtures, the bioluminescent signals of centrifugated LgBiT:SmBiT cell mixtures increased significantly. (**E**) Hela: NK-CD16 interactions were elevated by anti-EGFR antibody cetuximab. The bioluminescent signals were measured after 30-min incubation of Hela and NK-CD16, with or without cetuximab. Cetuximab elevated bioluminescent signals significantly in both E/T= 10 group and E/T= 5 group. Cetuximab-specific signals were higher in E/T= 10 group than E/T= 5 group. From (A) to (E), each data point represents one technical replicate. Error bars represent SD values. At least three independent biologic replicates were performed per experiment. MEM = membranous; cyto = cytosolic; E/T = effector/target cell ratio; WT = wild type.

The biosensor system was then characterized using different cell systems to determine the interaction probability between LgBiT and SmBiT. When co-expressed on the cell membrane, LgBiT-SmBiT bound and emitted strong luminescent signals (Figure 3C). In contrast, LgBiT-or SmBiT-positive cells alone had negligible background signals. When SmBiT and LgBiT expressed on the same cells (without spatial restriction), strong bioluminescence signals were detected. However, when SmBiT and LgBiT were expressed on separate cell membranes (SmBiT+ and LgBiT+ HeLa cells), the bioluminescent signal was low when both cell types were in suspension (Figure 3D). The intercellular interaction probability was increased by centrifugation, which resulted in 5-10 times higher bioluminescent signals in centrifuged LgBiT:SmBiT cell mixtures, indicating that the detecting signal is intercellular contact-dependent (Figure 3D).

The biosensor system was then applied to investigate antibody-induced cell-cell interactions. Specifically, intercellular interaction between NK-CD16 and Hela cells induced by an anti-EGFR antibody cetuximab was first evaluated. SmBiT^+^ NK-CD16 cells were incubated with LgBiT^+^ HeLa cells at different E:T ratios (5 or 10), with or without cetuximab (Figure 3E). Without cetuximab, no significantly different bioluminescent signals were detected in the effector-target cell mixtures regardless of E:T ratio (p = 0.5). The increased bioluminescent signals was observed in the presence of cetuximab, suggesting that the cetuximab promotes the interaction between NK-CD16 and HeLa cells. The bioluminescent signals were significantly higher in the 10 E:T ratio group than the group with 5 E:T ratio (p = 0.02), indicating cell density dependency.

### Interactions between NK and target cells induced by rituximab in a 3D system

NK-CD16 cells transfected with SmBiT with different linker lengths (SmBiT-0L, SmBiT-1L, SmBiT-3L, and SmBiT-12L cells) had comparable CD16 and HA expression levels (Figure 4A). All four SmBiT^+^NK-CD16 cell lines (SmBiT-0L, SmBiT-1L, SmBiT-3L, and SmBiT-12L) had substantial SmBiT expressions (Figure 4B). No significant difference in SmBiT expressions among the four SmBiT^+^NK-CD16 cell lines was observed (p = 0.1, one-way ANOVA). CD20 expression levels were largely comparable among CD20^+^LgBiT^+^ HeLa cell lines (LgBiT-0L, LgBiT-1L, LgBiT-3L, and LgBiT-12L cells) (Figure 4C). HeLa cells had relatively lower expressions of LgBiT-0L and LgBiT-1L and the expressions of LgBiT-3L and LgBiT-12L were comparable (Figure 4C).

**Figure 4.**
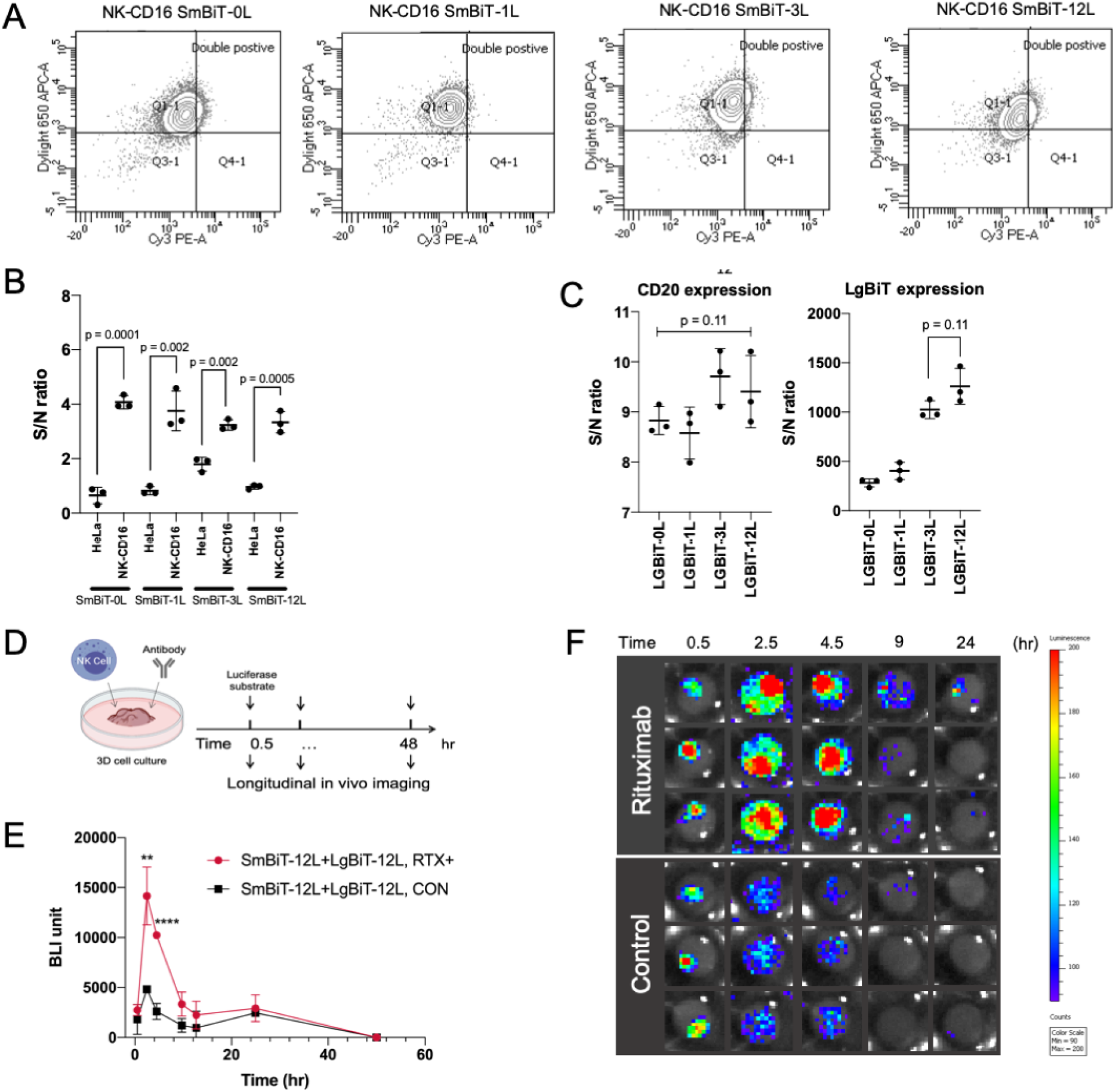
Constructing biosensor system in NK-CD16 and CD20+HeLa cells. (**A**) NK-CD16 cells stably expressing SmBiT (SmBiT-0L, SmBiT-1L, SmBiT-3L, and SmBiT-12L) were sorted by HA tag (Dylight 650) and CD16 (Cy3). (**B**) SmBiT expression in SmBiT-0L, SmBiT-1L, SmBiT-3L, and SmBiT-12L were comparable (p = 0.1, ordinary one-way ANOVA) and significantly higher than positive controls, i.e., HeLa cells transiently-expressing SmBiT. (**C**) CD20 expressions were comparable between CD20+LgBiT+ HeLa cell lines (LgBiT-0L, LgBiT-1L, LgBiT-3L, and LgBiT-12L cells, p = 0.11, ordinary one-way ANOVA). (D) LgBiT expression in CD20+LgBiT+ HeLa cell lines. No significant difference was observed in LgBiT expression in LgBiT-3L and LgBiT-12L cells. p = 0.11). In (B) and (C), each data point represents one technical replicate. Error bars represent SD values. At least three independent biologic replicates were performed per experiment. (**D**) Study design and bioluminescent images of NK-CD16 and CD20+HeLa interactions at 0.5, 2.5, 4.5, 9, and 24 hour. RTX = rituximab; CON = control. (**E**) (**F**) Bioluminescent intensity revealing direct interaction between NK-CD16 and CD20+HeLa cells. LgBiT-12L: SmBiT-12L pair was used. Each data point represents one technical replicate. Error bars represent SD values. At least three independent biologic replicates were performed per experiment.

The rituximab-induced NK-CD16 and CD20^+^HeLa interactions were assessed using LgBiT-12L: SmBiT-12L pair in the 3D cell culture system for 48 hours (Figure 4D-4F). Figure 4E and 4F showed the real-time NK-CD16 and CD20^+^HeLa interactions. The interactions were relatively low at the beginning of the imaging study (< 0.5 hr), then increased and peaked around the 2.5 hours, suggesting increasing frequency of effector and target cell interactions over time. Interestingly, the control group without rituximab also showed bioluminescent signals, indicating that NK-CD16 cells may interact with target cells mediated by other receptors such as IL2R^15, 16^. The interactions mediated by rituximab decreased over time from 2.5 to 9 hour.

Interactions gradually decreased after peaking, and were no longer detected at around 48 hours, which is consistent with the typical duration of antibody-dependent cell-mediated cytotoxicity (ADCC). At earlier time points (< 0.5 hr) the bioluminescent signals were relatively higher in the periphery than the central region, indicating that the interactions started in the periphery of the spheroids (Figure 4E and 4F).

### The optimized biosensor system with improved sensitivity

The biosensor pair was tested with varying combined spacer lengths and the rituximab-specific signals were normalized by the peak signals from the control group. The biosensor pair with the shortest theoretical combined spacer length (LgBiT-0L+ SmBiT-0L) had the lowest rituximab-specific signal, while the pair with the longest theoretical combined spacer length (LgBiT-12L: SmBiT-12L) had the highest rituximab-specific signal (Figure 5A). The results were also shown in quantitative data (Figure 5B) and longitudinal imaging profiles (Figure 5C).

**Figure 5.**
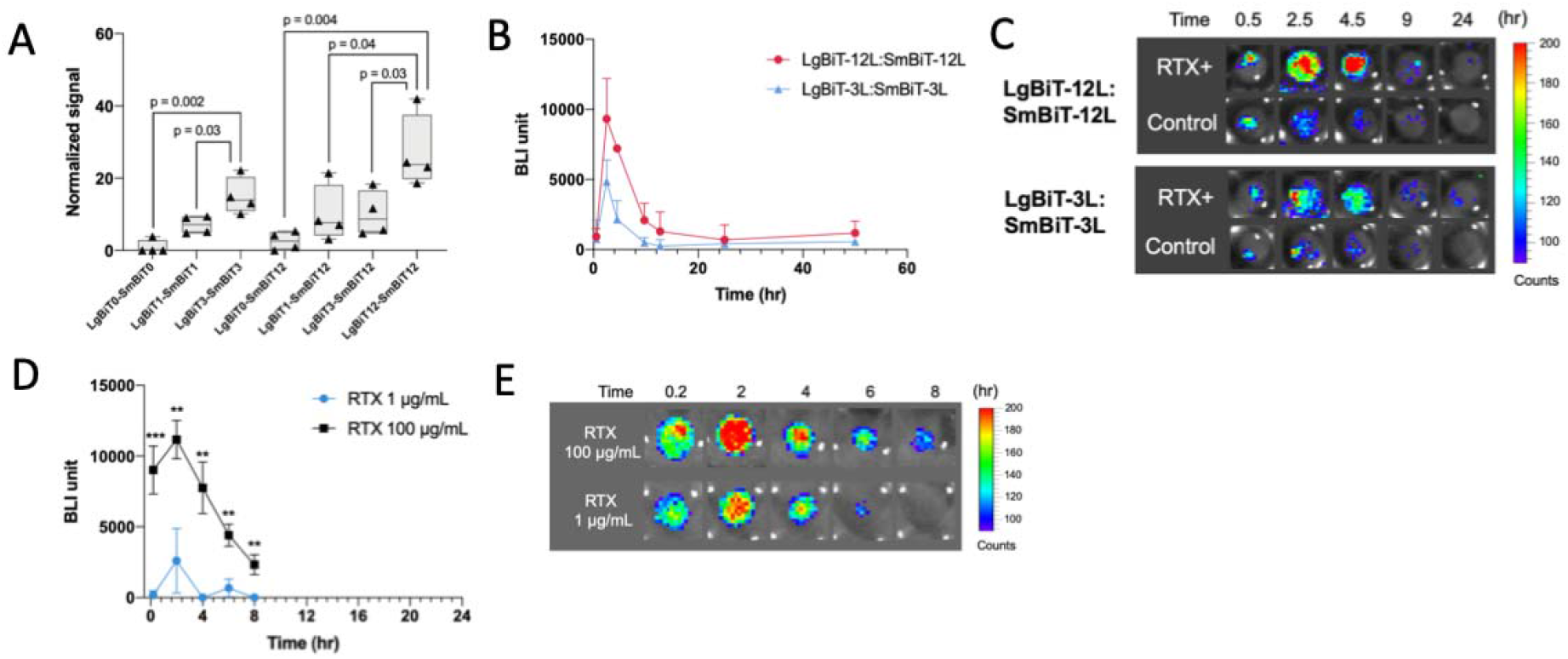
The optimization of sensitivity and selectivity of the biosensor system in the 3D cell culture and antibody dose dependency. (**A**) LgBiT-12L: SmBiT-12L pair showed the optimal sensitivity compared to other pairs. LgBiT-3L: SmBiT-3L pair showed higher normalized rituximab-specific signals than LgBiT-0L: SmBiT-0L and LgBiT-1L: SmBiT-1L pairs (p = 0.002 and p = 0.03). LgBiT-12L: SmBiT-12L pair showed higher normalized rituximab-specific signals than LgBiT-0L: SmBiT-12L, LgBiT-1L: SmBiT-12L, and LgBiT-3L: SmBiT-12L pairs (p = 0.004, p = 0.04, and p = 0.03). (**B**) Corrected bioluminescent signals indicated LgBiT-12L: SmBiT-12L pair showed higher selectivity than LgBiT-3L: SmBiT-3L pair. In (A) and (B), each data point represents one technical replicate. Error bars represent SD values. At least three independent biologic replicates were performed per experiment. (**C**) Bioluminescent images of NK-CD16 and CD20^+^HeLa interactions at 0.5, 2.5, 4.5, 9, and 24 hour. (**D**) Rituximab-induced effector: target cell clustering was antibody concentration-dependent. Raw data of bioluminescent signals from NK-CD16 and CD20^+^HeLa interactions, which was visualized by LgBiT-12L: SmBiT-12L pair. Each data point represents one technical replicate. Error bars represent SD values. At least three independent biologic replicates were performed per experiment. (**E**) Bioluminescent images of NK-CD16 and CD20^+^HeLa interactions at 0.2, 2, 4, 6, and 8 hour. RTX = rituximab.

Despite the rituximab-specific signals increased as the combined spacer lengths increased from 0 to 12 nm, the combined spacer length was not the only factor influencing the rituximab-specific signals selectivity, as the LgBiT-0L: SmBiT-12L pair had much lower rituximab-specific signals than the LgBiT-3L: SmBiT-3L pair (p = 0.005) (Figure 5A), suggesting lower detecting selectivity. Biosensor pairs with different combined spacer lengths, e.g., LgBiT-3L: SmBiT-3L, LgBiT-1L: SmBiT-12L, and LgBiT-3L: SmBiT-12L, had comparable rituximab-specific signals (p = 0.9). Based on these results, LgBiT-12L: SmBiT-12L biosensor pair was selected for subsequent dose-dependent study.

The intercellular interaction induced by rituximab was investigated at two concentrations (1 and 100 μg/mL). It was observed that the antibody-induced signal peaked at 2 hours in both dose groups, similar to previous observations. However, the high dose group (100 μg/mL) showed significantly higher cell-cell interaction throughout the imaging window compared to the low dose group, suggesting that there was an antibody concentration-dependent effect on the cell-cell interactions. This information is presented in Figure 5D and 5E, where the higher dose group shows a substantially higher level of cell-cell interaction compared to the low dose group.

### Interactions between T and tumor cells induced by blinatumomab

The intercellular interactions induced by the bispecific T cell engaging antibody blinatumomab were also assessed using the system (Figure 6A). The system sensitivity and selectivity were tested by comparing the signals across a range of linker lengths. Interestingly, the optimal linker pair was found to be LgBiT-3L: SmBiT-3L, which is shorter than the optimal length for NK and target cell interactions induced by rituximab, indicating cell line-specific optimal linkers. This finding is consistent with previous observations that natural killer (Dixon et al.) cells are capable of exerting their lytic effect at a relatively broader cell-cell distance compared to cytotoxic T cells (Chauveau, Aucher, Eissmann, Vivier & Davis, 2010). Figure 6B shows that the blinatumomab-induced T and target cell interactions peaked at 6 hrs, which is slower than the rituximab-induced NK and target cell interactions, suggesting different patterns of cell-cell interaction.

**Figure 6.**
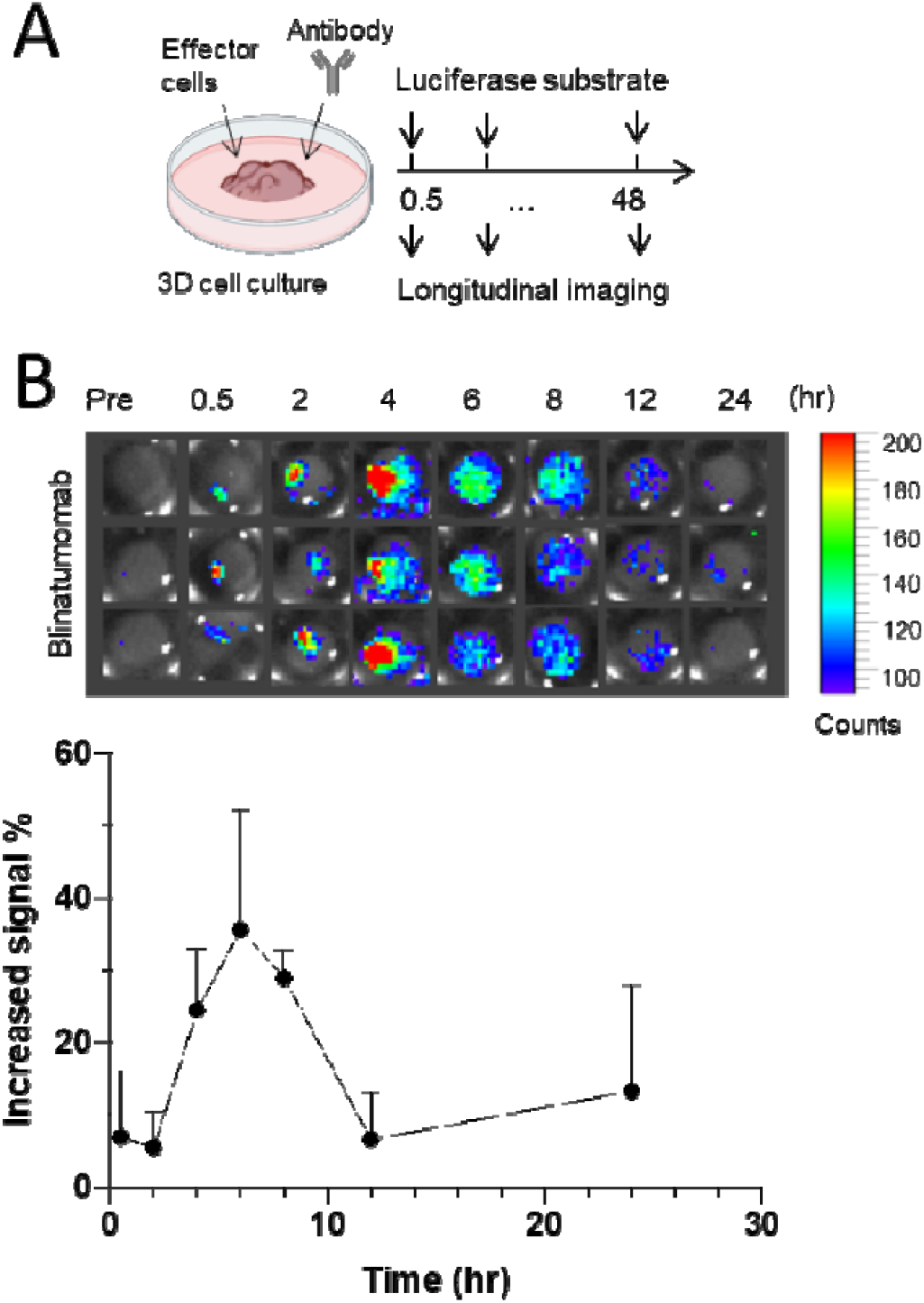
Interactions between T and target cells induced by blinatumomab. Raw data of bioluminescent signals from Jurkat T (CD3+) and Raj B (CD19) cell-cell interactions in the 3D cell culture system, which was visualized by LgBiT-3L: SmBiT-3L pair. Each data point represents one technical replicate. Error bars represent SD values. At least three independent biologic replicates were performed per experiment. (B) Bioluminescent images of at 0.2, 2, 4, 6, and 8 hour at 1 ng/ml blinatumomab.

## Discussion

The biosensor system described in this study provides a potential tool to address this need by enabling the visualization and quantification of intercellular interactions between immune cells and target cells. Specifically, it allows for the real-time monitoring of interactions between immune cells and target cells induced by therapeutic antibodies with high spatial and temporal resolution, which can help to uncover the underlying dynamics and assess its contribution to antibody treatment efficacy. Specifically, the biosensor system can be used to evaluate the interactions induced by different therapeutic antibodies, involving interactions between NK and tumor cells induced by rituximab and T and tumor cells induced by bispecific T cell engager. Overall, the biosensor system represents a promising platform for studying antibody-induced cell-cell interaction, which has potential applications in the development and optimization of antibody-based cancer therapies.

The use of split luciferase subunits as a biosensor is a powerful tool to detect cell-cell interactions and monitor the clustering dynamics between NK and target cells during ADCC. By equipping the split luciferases with different lengths of spacers, the biosensor system can accurately capture the distances between the opposing membranes in the NK-target cell synapse, allowing for improved sensitivity and selectivity in detecting cell contacts. The use of -(GGGGS)n-as the spacer confers flexibility and linearity to the biosensor system, contributing to its effectiveness in capturing elevated signals from antibody-induced NK-target cell interactions.

We observed that the sensitivity of the system did not solely depend on linker length. The biosensor pairs with inflexible split luciferase components (i.e., no spacer, LgBiT-0L) had the lowest S/N ratios regardless of the theoretical combined spacer lengths, suggesting that certain degrees of flexibility of the biosensor system are required for interactions. Yielding approximately 12, 26, and 30 nm theoretical combined spacer lengths (LgBiT-3L: SmBiT-3L, LgBiT-1L: SmBiT-12L, and LgBiT-3L: SmBiT-12L, respectively), which are close to the theoretical distances between opposing cell membranes in a synapse (15 – 40 nm) (McCann et al., 2003), the biosensor pairs showed similar rituximab-specific bioluminescent signals. The pair with longer theoretical combined spacer lengths did not offer superior sensitivity (Figure 5A). These observations could possibly be explained by the unstable distances between the opposing cell membranes in synapses. Rather than rigid surfaces, cell membranes in the synapse are constantly moving and changing along with synapse formation progress. The weak binding (190 μM) between the luciferase subunits in the biosensor system can be easily interrupted by the membrane movement even though the biosensor pairs’ theoretical combined spacer lengths were adequate. LgBiT-12L: SmBiT-12L pair provided the highest sensitivity in detecting cell conjugations, possibly due to the proper flexibility on both sides.

The biosensor system allowed for real-time monitoring of NK: target cell interactions and clustering dynamics, providing spatial and temporal resolution. ADCC, mediated by NK cells, is a crucial mechanism of action for many therapeutic antibodies, such as tranzusumab, rituximab, cetuximab, and antibodies that trigger stronger effector functions are increasingly being developed (Wang & Weiner, 2008). ADCC is a dynamic process that is influenced by multiple factors, such as the concentration and distribution of the antibody and effector cells, the tumor microenvironment, and the expression of Fc receptors on the effector cells (Barreira da Silva, Graf & Munz, 2011; Cartron et al., 2002; Chang et al., 2021; Helguera et al., 2011; Musolino et al., 2008). However, the mechanisms and dynamics of ADCC in vivo and the contribution of ADCC to antibody treatment responses have not been completely understood; most measurements of ADCC consist of bulk assays or discontinuous methods, obscuring the spatial and temporal resolutions of antibody-mediated cell-cell interactions and lacking the physiological context. The biosensor system we developed provides a powerful tool for noninvasively assessing ADCC in a 3D culture system, enabling the investigation of the mechanisms and dynamics of ADCC at the population levels with sufficient spatial and temporal resolutions. The biosensor system was also able to detect the concentration-dependent effects of antibodies on NK: target cell interactions, which could be extended to investigate the dynamics of ADCC in vivo and in different disease models, facilitating the development and optimization of antibody-based therapies.

The system often temporal characterizations of cell-cell interactions induced by antibodies. Interestingly, brief bioluminescent signals were detected in the control without rituximab, suggesting that there may be other receptors, such as IL2R, involved in mediating NK cell interactions with target cells (Chen, Kawashima, Lowe, Lanier & Fukuda, 2005; Orange, 2008). The rituximab group showed significantly higher peak signals compared to the control group, indicating that rituximab increased NK: target cell conjugation frequencies. The peak time of cell conjugation was around 2 hours, possibly resulting from rituximab and NK cell distribution kinetics or sequential killing effects of NK cells. Similar findings were observed in the cell conjugation kinetics at different rituximab concentrations. These findings agreed with previous studies regarding the time scale of antibody-medicated cellular cytotoxicity (Kamen, Thakurta, Myneni, Zheng & Chung, 2019; Li et al., 2010). While rituximab can opsonize target cells rapidly, the slower trafficking of NK cells could delay the establishment of the first cell conjugate, which was around 80 minutes for most NK cells in a previous study (Li et al., 2010). Of note, blinatumomab-induced T and target cell interactions peaked at 6 hours, which is slower than the peak observed in the rituximab-induced NK and target cell interactions. T cells are often referred to as serial killers because they can recognize and kill multiple target cells in a sequential manner, as opposed to NK cells, which can rapidly kill target cells without prior sensitization.

The slower peak observed in the blinatumomab-induced T and target cell interactions may be due to the need for T cells to be extensively activated, leading to a slower onset of target cell killing compared to the more rapid and innate killing ability of NK cells. However, it is important to note that the specific mechanisms of action and experimental conditions for each drug may also play a role in the observed differences in cell-cell interaction kinetics, which warrant further investigation.

Overall, the proximity-based biosensor system described in the study provides a promising tool for noninvasive and continuous monitoring of cell clustering dynamics induced by various antibody therapies, including canonical antibodies and bispecific T-cell engagers. However, further investigation is needed to fully understand the complex physiological factors and establish the relationship between cell conjugation and cell-killing effects in vivo. The limitations of the study suggest areas for future research, which could potentially lead to the development of more effective cancer treatments.

## Author Contributions

Conceptualizations: Y.T., and Y.C.; methodology: Y.T., and Y.C.; formal analysis: Y.T., and Y.C.; investigation: J Y.T., and Y.C.; writing-original draft: Y.T., and Y.C.; writing-reviewing and editing: Y.T., and Y.C.; supervision: Y.C.

## References

Armingol E, Officer A, Harismendy O, & Lewis NE (2021). Deciphering cell-cell interactions and communication from gene expression. Nat Rev Genet 22: 71–88.

Baginska J, Viry E, Paggetti J, Medves S, Berchem G, Moussay E, et al. (2013). The critical role of the tumor microenvironment in shaping natural killer cell-mediated anti-tumor immunity. Front Immunol 4: 490.

Barreira da Silva R, Graf C, & Munz C (2011). Cytoskeletal stabilization of inhibitory interactions in immunologic synapses of mature human dendritic cells with natural killer cells. Blood 118: 6487–6498.

Bechtel TJ, Reyes-Robles T, Fadeyi OO, & Oslund RC (2021). Strategies for monitoring cell-cell interactions. Nat Chem Biol 17: 641–652.

Cartron G, Dacheux L, Salles G, Solal-Celigny P, Bardos P, Colombat P, et al. (2002). Therapeutic activity of humanized anti-CD20 monoclonal antibody and polymorphism in IgG Fc receptor FcgammaRIIIa gene. Blood 99: 754–758.

Chang HY, Wu S, Li Y, Zhang W, Burrell M, Webster CI, et al. (2021). Brain pharmacokinetics of anti-transferrin receptor antibody affinity variants in rats determined using microdialysis. MAbs 13: 1874121.

Chauveau A, Aucher A, Eissmann P, Vivier E, & Davis DM (2010). Membrane nanotubes facilitate long-distance interactions between natural killer cells and target cells. Proc Natl Acad Sci U S A 107: 5545–5550.

Chen S, Kawashima H, Lowe JB, Lanier LL, & Fukuda M (2005). Suppression of tumor formation in lymph nodes by L-selectin-mediated natural killer cell recruitment. J Exp Med 202: 1679–1689.

Dixon AS, Schwinn MK, Hall MP, Zimmerman K, Otto P, Lubben TH, et al. (2016). NanoLuc Complementation Reporter Optimized for Accurate Measurement of Protein Interactions in Cells. ACS Chem Biol 11: 400–408.

Dustin ML (2014a). The immunological synapse. Cancer Immunol Res 2: 1023–1033.

Dustin ML (2014b). What counts in the immunological synapse? Mol Cell 54: 255–262.

Helguera G, Rodriguez JA, Luria-Perez R, Henery S, Catterton P, Bregni C, et al. (2011). Visualization and quantification of cytotoxicity mediated by antibodies using imaging flow cytometry. J Immunol Methods 368: 54–63.

Kamen L, Thakurta T, Myneni S, Zheng K, & Chung S (2019). Development of a kinetic antibody-dependent cellular cytotoxicity assay. J Immunol Methods 468: 49–54.

Li G, Zhang L, Chen E, Wang J, Jiang X, Chen JH, et al. (2010). Dual functional monoclonal antibody PF-04605412 targets integrin alpha5beta1 and elicits potent antibody-dependent cellular cytotoxicity. Cancer Res 70: 10243–10254.

McCann FE, Vanherberghen B, Eleme K, Carlin LM, Newsam RJ, Goulding D, et al. (2003). The size of the synaptic cleft and distinct distributions of filamentous actin, ezrin, CD43, and CD45 at activating and inhibitory human NK cell immune synapses. J Immunol 170: 2862–2870.

Miah SM, & Campbell KS (2010). Expression of cDNAs in human Natural Killer cell lines by retroviral transduction. Methods Mol Biol 612: 199–208.

Musolino A, Naldi N, Bortesi B, Pezzuolo D, Capelletti M, Missale G, et al. (2008). Immunoglobulin G fragment C receptor polymorphisms and clinical efficacy of trastuzumab-based therapy in patients with HER-2/neu-positive metastatic breast cancer. J Clin Oncol 26: 1789–1796.

Nishida-Aoki N, & Gujral TS (2019). Emerging approaches to study cell-cell interactions in tumor microenvironment. Oncotarget 10: 785–797.

Orange JS (2008). Formation and function of the lytic NK-cell immunological synapse. Nat Rev Immunol 8: 713–725.

Redman JM, Hill EM, AlDeghaither D, & Weiner LM (2015). Mechanisms of action of therapeutic antibodies for cancer. Mol Immunol 67: 28–45.

Shelton SE, Nguyen HT, Barbie DA, & Kamm RD (2021). Engineering approaches for studying immune-tumor cell interactions and immunotherapy. iScience 24: 101985.

Tang Y, & Cao Y (2021). Modeling Pharmacokinetics and Pharmacodynamics of Therapeutic Antibodies: Progress, Challenges, and Future Directions. Pharmaceutics 13.

Waldman AD, Fritz JM, & Lenardo MJ (2020). A guide to cancer immunotherapy: from T cell basic science to clinical practice. Nat Rev Immunol 20: 651–668.

Wang SY, & Weiner G (2008). Complement and cellular cytotoxicity in antibody therapy of cancer. Expert Opin Biol Ther 8: 759–768.

Yang J, Zhang Z, Lin J, Lu J, Liu BF, Zeng S, et al. (2007). Detection of MMP activity in living cells by a genetically encoded surface-displayed FRET sensor. Biochim Biophys Acta 1773: 400–407.

Zhang D, He W, Wu C, Tan Y, He Y, Xu B, et al. (2019). Scoring System for Tumor-Infiltrating Lymphocytes and Its Prognostic Value for Gastric Cancer. Front Immunol 10: 71.

